# Refined mechanism of promoter Nucleosome-Depleted Regions resetting after replication

**DOI:** 10.1101/2024.04.14.589417

**Authors:** Sevil Zencir, Jatinder Kaur Gill, Françoise Stutz, Julien Soudet

## Abstract

Replication disrupts chromatin organization. Thus, the rapid resetting of nucleosome positioning is essential to maintain faithful gene expression. The initial step of this reconfiguration occurs at Nucleosome-Depleted Regions (NDRs). While studies have elucidated the role of Transcription Factors (TFs) and Chromatin Remodelers (CRs) *in vitro* or in maintaining NDRs *in vivo*, none has addressed their *in vivo* function shortly after replication. Through purification of nascent chromatin in yeast, we dissected the choreography of events governing the proper positioning of the −1/+1 nucleosomes flanking promoter NDRs. Our findings reveal that CRs are the primary contributors of −1/+1 repositioning post-replication, with RSC acting upstream of INO80. Surprisingly, while Reb1 and Abf1 TFs are not essential for NDR resetting, they are required for NDR maintenance *via* the promotion of H3 acetylations. Altogether, we propose a two-step model for NDR resetting in *S. cerevisiae*: first, CRs alone reset promoter NDRs after replication, while a combination of TFs and CRs is required for subsequent maintenance.

**Teaser:** RSC acts upstream of INO80 for NDR re-establishment after replication followed by a combined action of CRs and TFs for NDR maintenance.

## INTRODUCTION

Fundamental eukaryotic cellular processes such as transcription and replication are intricately connected to chromatin organization [1]. Nucleosomes constitute the fundamental building blocks of eukaryotic chromatin. They are composed of a tetramer called (H3-H4)_2_, which is surrounded by two dimers of H2A-H2B. 147 base pairs (bp) of genomic DNA makes 1.65 turns around this organizing unit [2]. Each of the histone proteins can undergo post-translational modifications such as acetylation, methylation and many others that will affect DNA processing [3]. The main function of nucleosomes is certainly to hinder spurious DNA-based mechanisms by restricting the access of DNA-binding proteins to the DNA [4]. Consequently, eukaryotic genomes consist in arrays of closely spaced nucleosomes separated by Nucleosome-Depleted Regions (NDRs) which constitute the starting point of DNA-related mechanisms such as transcription and replication [1]. However, nucleosomes are disrupted ahead of the replication fork in S-phase and nascent chromatin needs to be rapidly reset in order to maintain a faithful pattern of gene expression [5, 6]. Thus, the mechanism of NDR restoration after replication appears as the first step to preserve a proper transcriptional program.

Promoter NDRs are at the heart of the transcription initiation process since they provide access to the Transcription Factors (TFs), cofactors (coactivators, corepressors and chromatin-modifying enzymes) and the RNA polymerase II (RNAPII) machinery [7]. They are flanked by the so-called −1 and +1 nucleosomes, the latter being the closest to the Transcription Start Site (TSS) of coding genes [1, 8]. They can be quite heterogenous in size within, as well as among species [9]. However, for expressed genes, their overall average size is relatively constant from lower eukaryotes to multicellular organisms with a distance of 300 bp between the −1 and +1 dyads. In *Saccharomyces cerevisiae* (*S. cerevisiae*), NDRs are featured by poly-dA/dT sequences that intrinsically promote nucleosome depletion [10–13]. These elements often surround TFs specific motifs. A TATA or TATA-like motif can be found at an average distance of 100 bp upstream of the +1 dyad, implying that a 25 bp left-shift of the +1 nucleosome might hinder the recruitment of the transcription initiation machinery [14–16].

One important class of protein complexes which functions in setting a proper distance between −1/+1 nucleosomes are the ATP-dependent Chromatin Remodelers (CRs) [17, 18]. By using the energy delivered *via* ATP hydrolysis, remodelers alter DNA-histone interactions resulting in sliding and/or eviction of nucleosomes. Two main CRs are involved in setting NDRs in *S. cerevisiae*, the SWI/SNF-related RSC (Remodeling the Structure of Chromatin), related to the PBAF CR in mammals, and the INO80 complex, conserved among eukaryotes [17]. *In vitro*, incubation of RSC with yeast reconstituted chromatin produces over-extended NDRs at promoters compared to *in vivo* [19, 20]. This activity is stimulated by promoter poly-dA/poly-dT tracts [14, 19, 21]. In contrast, INO80 sets the proper positioning of the +1 nucleosome of poly-dA/poly-dT enriched promoters [19, 22]. *In vivo*, RSC depletion leads to a left and right shift of the +1 and −1 nucleosomes, respectively, while INO80 removal induces an opposite effect [14, 23, 24]. Thus, RSC is considered as a “pusher” while INO80 appears as a “puller” setting in conjunction the accurate placement of −1/+1 nucleosomes.

Another class of factors known to be involved in NDR width setting is a subset of TFs known as Nucleosome-Displacing Factors (NDFs) [25, 26]. *In vitro*, Reb1 and Abf1 act as barrier factors bound to specific DNA sequence motifs that coordinate, together with several CRs, the positioning of nucleosome arrays [4, 19, 27]. In yeast, depletion or mutants of Abf1 and Reb1 lead to a decrease of NDR width for the subset carrying the corresponding DNA-binding motifs [14, 28–30].

Is there a temporal order between CRs and TFs/NDFs in resetting promoter NDRs? The current model mainly comes from data obtained in higher eukaryotes. Some TFs, named Pioneer factors (PFs), can interact with DNA-bound nucleosomes *in vitro* [31]. This contact promotes a partial unwrapping of the DNA around nucleosomes that subsequently initiate the action of CRs to form NDRs. In such a model, PFs are acting upstream of CRs to trigger chromatin opening. Interestingly, *in vitro*, yeast Reb1 targets its motif within nucleosomes inducing their partial unwrapping [32]. However, *in vivo* depletion of Reb1 and RSC is additive suggesting independent functions for Reb1 and RSC [14], questioning the potential generality of the PF model.

Several recent studies have tackled the maturation of chromatin following replication [6]. In yeast, they focused on the fast reestablishment of nucleosomes within gene bodies, highlighting the role of transcription and histone chaperones in this process [33–35]. Studies in Drosophila S2 cells have shown that resetting of NDRs is dependent on the reassociation of TFs while some data in mouse Embryonic Stem Cells (mESCs) indicate that the transcription process participates in restoring NDRs accessibility [36, 37]. However, none of these studies tested the absence of TFs or CRs in NDR resetting following replication. Through specific labeling of nascent chromatin coupled with nuclear depletion of TFs or CRs, we aimed to dissect the *in vivo* mechanism of promoter NDR resetting after replication in this study. We propose a temporally unexpected scenario in which CRs act upstream of TFs.

## RESULTS

### Purification of nascent chromatin after a short EdU pulse

In order to map the position of nucleosomes on the nascent DNA, we used an approach in which *S. cerevisiae* cells engineered for the uptake of 5-Ethynyl-2′-deoxyUridine (EdU), an analog of thymidine, are incubated in an EdU-containing medium for a short pulse [38]. Considering the rapid average speed of the replication fork in *S. cerevisiae* cells and the rapid maturation of yeast chromatin [34, 39], we opted for a 2 minute (min) DNA labelling. As a proof of principle, we assayed whether a 2 min pulse was enough to specifically detect newly replicated DNA at early replication origins as compared to the late ones in an S-phase synchronized population of cells (fig. S1A). To perform this, we synchronized *S. cerevisiae* cells in G1-phase before releasing them into S-phase for 30 min in order to reveal the early Autonomously Replicated Sequences (ARSs). After 30 min, EdU was added for 2 min in both G1 arrest and during S-phase release. The chromatin was then extracted, MNase digested to obtain the footprint of nucleosomes along the DNA, labelled with biotin *via* a Click-it reaction and purified by streptavidin beads. Finally, the eluted material was paired-end sequenced (fig. S1A). Indeed, the purification of 2 min EdU-labelled DNA was specific enough to reveal the early ARSs (fig. S1B and C). Thus, despite a short pulse of EdU labelling, our experimental set-up allows the detection of newly replicated chromatin among a large majority of non-labelled DNA.

### Reb1 and Abf1 TFs are dispensable for −1/+1 positioning shortly after replication

After assessing its validity, we implemented our experimental scheme in anchor-away strains in which either TFs, CRs or RNAPII can be conditionally depleted by Rapamycin (Rap) treatment (Fig. 1A). To assess the position of virtually all the nucleosomes on newly replicated chromatin and to not restrict our analyses to early replicated areas, we performed the experiments with non-synchronized cells. We also paired-end sequenced the mature chromatin in parallel in order to compare the positioning of steady-state nucleosomes with their respective positioning in nascent chromatin.

**Fig. 1:**
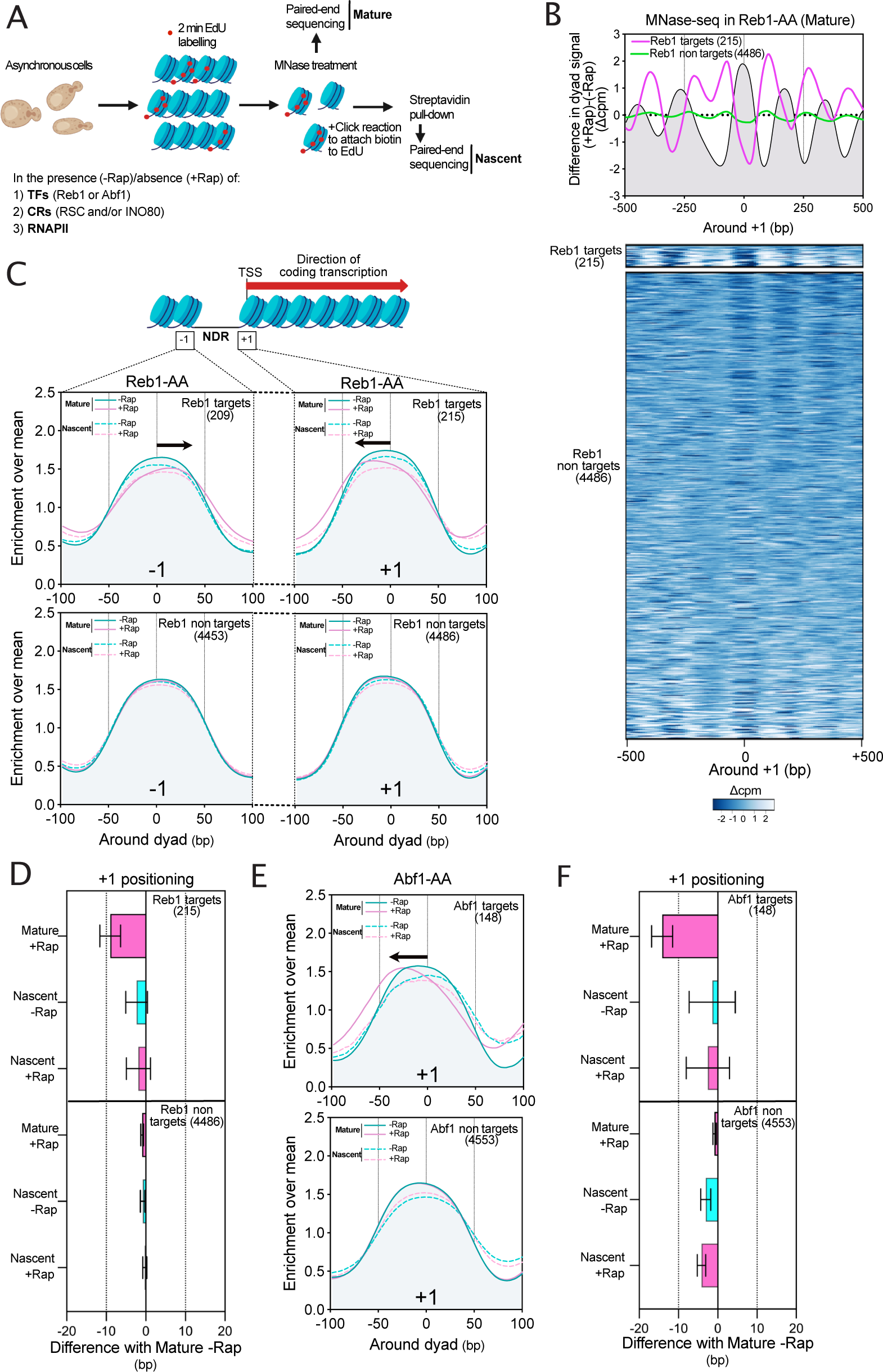
Reb1 and Abf1 Transcription Factors (TFs) are dispensable for proper −1/+1 nucleosomes positioning after replication. (**A**) Schematic representation of the experimental design for EdU labeling and MNase-seq: asynchronous *S. cerevisiae* cells engineered for EdU uptake were treated or not with Rapamycin (Rap) for 30min in order to deplete either Transcription factors (TFs), Chromatin remodelers (CRs) or RNA Polymerase II (RNAPII). EdU was then added in the medium for a pulse of 2min. Cells were crosslinked with formaldehyde, the DNA was extracted and treated with MNase. The click reaction was then performed for attaching biotin to EdU-labeled DNA and followed by a streptavidin pull-down. After library preparation, the purified DNA was paired-end sequenced giving the information of nucleosome positioning on ‘Nascent’ chromatin. In parallel, MNase-digested DNA was aliquoted before click reaction and paired-end sequenced to extract the nucleosome positioning of ‘Mature’ chromatin. (**B**) Top panel: Metagene plot showing the difference of nucleosome positioning in +Rap and -Rap conditions for a Reb1-AA strain. Genes were divided into two groups as ‘Reb1 targets’ and ‘Reb1 non targets’, the number of genes in each cluster is indicated in brackets. This plot is centered on the +1 dyad. Bottom panel: Heatmap of the difference in MNase-seq for the two classes of genes. (**C**) Metagene analysis showing the mean positioning of −1 and +1 nucleosomes (left and right panels, respectively) for the two classes of genes for both Mature and Nascent chromatin in the presence (-Rap) or absence (+Rap) of Reb1. Data were represented with an enrichment over mean method described in Materials and Methods. (**D**) Bar plot showing the distribution of the difference of +1 peaks in a given condition relative to the Mature -Rap position for the Reb1-AA strain. The bar represents the mean of the genes in different groups and the error bars represent the 95% confidence intervals. (**E**) Same as in C for the +1 nucleosome in the presence (-Rap) or absence (+Rap) of Abf1. (**F**) Same as in D but illustrating the results of Abf1-AA.

We first tested the importance of Reb1 TF in resetting promoter NDR architecture after replication. Reb1 depletion only affects the expression of a minority of genes that was previously defined [40, 41]. Thus, we performed our analyses by making the distinction between Reb1 ‘target’ and ‘non-target’ genes. As already reported, Reb1 depletion leads to a shrinkage of the promoter NDR on the mature chromatin at the target genes (Fig. 1B) [14, 28]. We then focused on the positioning of the +1 nucleosome (Fig. 1C, right panel and fig. S1D).

Comparison of nascent and mature +1 positioning in the absence of Rap for the non-target genes revealed that within 2min of EdU, most of the NDR-flanking nucleosomes were correctly positioned in agreement with a rapid resetting of promoter NDRs in *S. cerevisiae* (Fig. 1C and D, bottom panels) [34]. However, the positioning of the +1 nucleosome was significantly different between nascent and mature chromatin for target genes in the absence of Reb1 (Fig. 1C and D, top panels and fig. S1D). Indeed, while mature +1 nucleosomes are shifted upstream, nascent +1 are positioned as in the -Rap condition. We made a similar observation for the −1 nucleosome, which is shifted downstream in the mature chromatin (Fig. 1C left).

We performed the same analysis by depleting another TF, Abf1. Genes were split into ‘targets’ and ‘non-targets’ (fig. S1E) [42]. +1 nucleosome positioning on nascent chromatin in the absence of Abf1 is not significantly different to its positioning in a -Rap condition while mature +1 is shifted upstream (Fig. 1E and F, upper panels). The −1 nucleosome shows the same trend (fig. S1F).

Taken together, these results indicate that Reb1 and Abf1 TFs are not required for establishing a proper positioning of the −1/+1 nucleosomes immediately after replication but participate in the long-term maintenance of their mature positioning.

### Maintenance of histone acetylation rescues −1/+1 positioning defects caused by the absence of Reb1

Our previous work proposed that H3K18 acetylation (H3K18ac) on +1 nucleosomes is associated with a right shift while H3K36 trimethylation (H3K36me3) is associated with a left shift [43]. We decided to deepen this observation by systematically assaying the positioning of modified +1 nucleosomes. To perform this, we reanalyzed the dataset generated by the Friedman and Rando labs [44]. In this work, the MNase-ChIP-seq of the different histone modifications generated an unprecedented high resolution of modified nucleosomes. We reasoned that the sequencing of the input would indicate the sum of all the unmodified/modified nucleosomes while the mapping of a given modification would indicate its specific positioning relative to this sum (Fig. 2A). Thus, we subtracted the corresponding input from its associated modifications and plotted the signal as a Z-score centered on the +1 nucleosome dyad (Fig. 2B). All H3 acetylated nucleosomes are right shifted as compared to the position of the +1 nucleosome dyad (Fig. 2B). Moreover, the H3K18ac profile obtained by MNase-ChIP-seq from our previous work shows a similar behavior [43]. These analyses happen to be true for the −1 nucleosome but instead with an associated left shift implying that histone H3 acetylation increases promoter NDR length (fig. S2A). While it has been known that acetylation of histone H4 positively correlated with gene expression, we observed that the acetylation of different lysine residues on histone H4 does not induce a similar shift in −1/+1 positioning (fig. S2C).

**Fig. 2:**
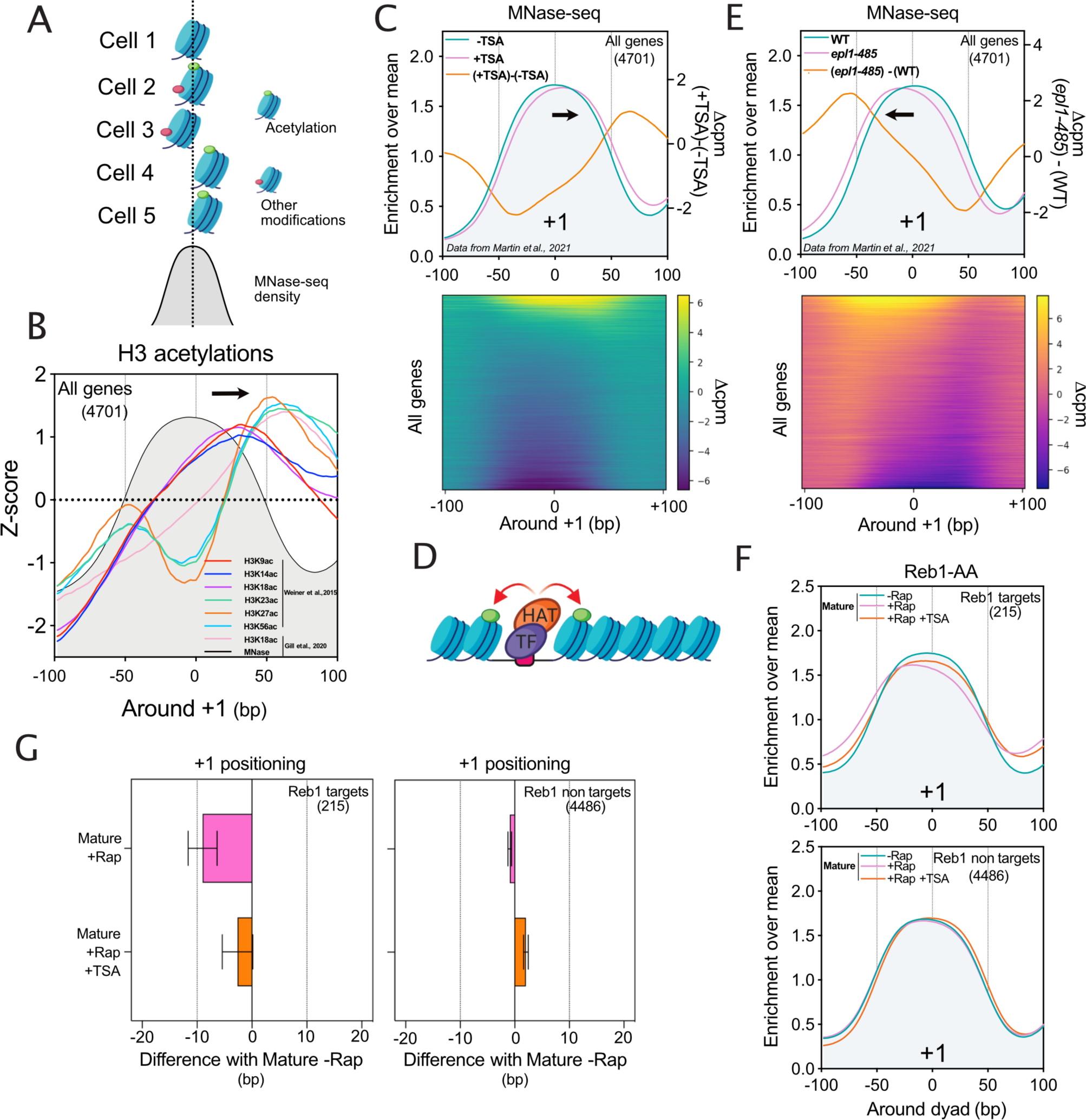
Effect of histone acetylation on −1/+1 nucleosome positioning. (**A**) Nucleosome positioning obtained by MNase-seq averages the positions of the differently modified nucleosomes. (**B**) Mean distribution of acetylation of different lysine residues on histone H3 around the +1 dyad. Reanalyzing data from Weiner et al., 2015 and Gill et al., 2020, the positioning of modified nucleosomes related to their respective total MNase-seq signal was assessed (see Materials and Methods). The mean of the difference between the histone H3 acetylation and its respective input is represented and normalized as a Z-score. (**C**) Top panel: Average metagene plot of +1 positioning obtained by MNase-seq from cells treated or not with TSA. Data were obtained from Martin et al., 2021. Bottom panel: Heatmap of the difference in MNase-seq signal of cells treated with TSA and without TSA for all genes (4701). (**D**) Scheme showing the role of TFs in the recruitment of HATs in order to acetylate nucleosomes flanking the NDR. (**E**) Top panel: Average metagene profile of +1 positioning obtained by MNase-seq from cells expressing wild-type *EPL1* or mutant *epl1-485*, a subunit of the NuA4 HAT complex. Data were obtained from Martin et al., 2021. Bottom panel: Heatmap of the difference in MNase-seq of cells expressing *EPL1* or *epl1-485* for all genes (4701). (**F**) Average metagene plot of MNase-seq signal in Reb1-AA treated or not with Rap and TSA. (**G**) Bar plot showing the distribution of the difference of +1 peaks in a given condition relative to the Mature -Rap position in Reb1-AA.

To confirm the concept that histone H3 acetylated +1 nucleosomes are right-shifted, we reanalyzed MNase-seq data of cells treated with Trichostatin A (TSA) in order to inhibit Histone deacetylases (HDACs) and to increase histone H3 acetylation levels [45]. Indeed, inhibition of deacetylation leads to a right shift of the +1 nucleosome and a left shift of the −1 nucleosome (Fig. 2C and fig. S2D).

Histone acetyltransferases (HATs) directly interact with transcriptional activators to mediate −1/+1 histone acetylations (Fig. 2D). Epl1 is a component of two HATs: NuA4 and Piccolo [46] complexes both feature a HAT module comprising Esa1, Epl1, and Yng2. Notably, only NuA4 contains Tra1, mediating the interaction of this HAT with transcription activators [47]. When the 348 C-terminal amino acids of Epl1 (Epl11–485) are deleted, it disrupts the incorporation of the HAT module into NuA4. As a result, cells expressing Epl11–485 are believed to have Piccolo but lack NuA4 [46]. We took advantage of published MNase-seq data using the *epl1-485* mutant [45]. This mutant reveals a left shift of most, if not all, of the +1 nucleosomes while the −1 nucleosomes are right shifted (Fig. 2E and fig. S2E), supporting that Epl1-dependent histone acetylation promotes NDR opening.

Finally, we tested the −1/+1 positioning in mature chromatin upon TSA treatment in the absence of Reb1 TF. Our analysis revealed that maintenance of histone acetylation is able to compensate for the lack of Reb1 (Fig. 2F and G and fig. S2F). Thus, Reb1 does not have a nucleosome displacing factor activity by itself but it is more certainly the Reb1-mediated acetylation that properly positions −1/+1 nucleosomes.

### The RSC ATPase activity and INO80 are essential for −1/+1 nucleosome repositioning after replication

RSC and INO80 are the two main CRs involved in NDR setting *in vitro* [19]. Moreover, *in vivo*, RSC has been described as a pusher while INO80 is acting as a puller [23]. These two CRs are targeting most, if not all, promoter NDRs in *S. cerevisiae*. Indeed, RSC depletion *via* nuclear depletion of the ATPase catalytic subunit Sth1 leads to a shrinkage of promoter NDRs over the whole genome (fig. S3A). Contrarily, nuclear depletion of Ino80 carrying the ATPase activity of INO80 globally induces an expansion of the promoter NDRs (fig. S3B).

Analysis of the +1 position on nascent chromatin in the absence of Sth1 revealed that, contrary to GRFs, the left shift is already visible on the newly synthesized chromatin (Fig. 3A and B and fig. S3E). Moreover, the right shift in the absence of Ino80 is also detected at the level of nascent chromatin (Fig. 3C and D and fig. S3F). The phenotype of the depletion of these two CRs appears shortly after replication indicating that they act upstream of TFs for promoter NDR resetting. Symmetrical results were obtained with the −1 nucleosome (fig. S3C and D).

**Fig. 3:**
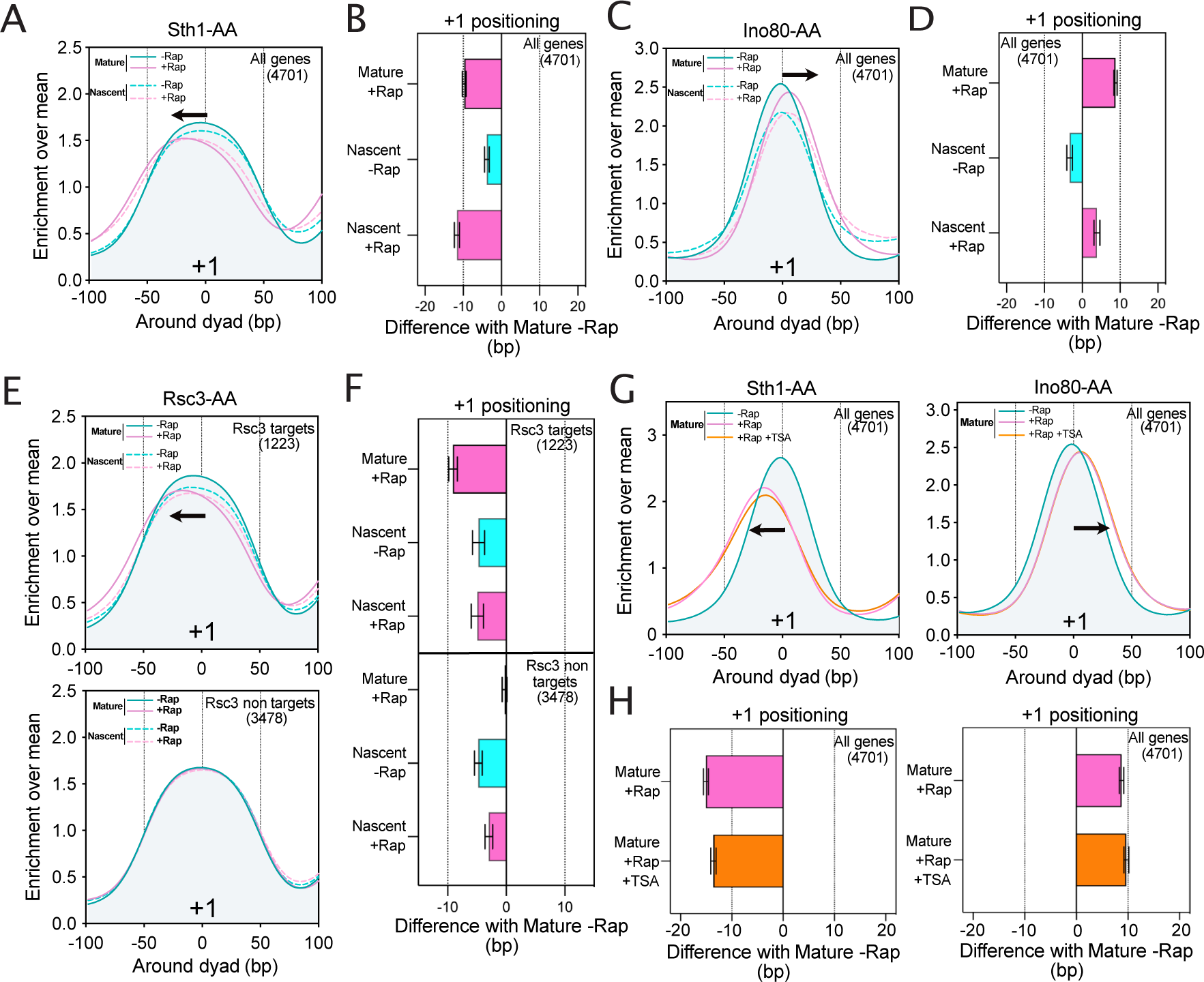
RSC and INO80 Chromatin Remodelers (CRs) rapidly position −1/+1 nucleosomes after replication. (**A**) Metagene plot showing the mean positioning of the +1 nucleosome for all genes for both ‘Mature’ and ‘Nascent’ chromatin in the presence or absence of Sth1. (**B**) Bar plot showing the distribution of the difference of +1 peaks in a given condition relative to the Mature -Rap position for the Sth1-AA strain. (**C**) Same as A, but showing the results for Ino80-AA. (**D**) Same as B, showing the distribution of the difference of +1 peaks for Ino80-AA. (**E**) Same as A, illustrating the results for Rsc3-AA. Genes were divided into two classes as ‘Rsc3 targets’ and ‘Rsc3 non targets’. (**F**) Same as B, for Rsc3-AA taking into consideration the two classes of genes. (**G**) Metagene profile of the mean +1 positioning in Sth1-AA and Ino80-AA for mature chromatin in -Rap, +Rap and +Rap +TSA conditions. (**H**) Same as B and D, but showing the results for Sth1- and Ino80-AA in the presence of TSA.

One proposed model for RSC recruitment to promoter NDRs is through the recognition of a CGSG motif on the DNA *via* the Rsc3/Rsc30 DNA interaction module [48, 49]. Surprisingly, depletion of the essential Rsc3 subunit of the RSC complex leads to the shrinkage of only a fraction of promoter NDRs as compared to the Sth1 nuclear depletion (fig. S3G), allowing us to define ‘target’ and ‘non-target’ genes in the absence of Rsc3. With respect to the chromatin maturation after replication, depletion of Rsc3 does not change the +1 or −1 positioning as compared to the -Rap condition (Fig. 3E and F and fig. S3H and I). This conveys that the Rsc3 DNA binding module is not required for the reestablishment of the +1 nucleosome after replication but is essential for its long-term correct placement as observed in the case of TFs depletion. In other words, the RSC ATPase and DNA-binding modules clearly show different activities regarding NDR formation and maintenance, respectively.

Finally, TSA treatment did not change the effect of Ino80 and Sth1 depletion on mature chromatin implying that CRs are acting upstream of histone acetylation (Fig. 3G and H).

### RSC ATPase activity is required for INO80 recruitment and puller activity

Although our data indicate that RSC and INO80 CRs are necessary for promoter NDR formation after replication, while TFs participate in a second step to their maintenance, whether a temporal hierarchy exists between RSC and INO80 remains to be clarified. To address this question, we performed the analysis of the +1 positioning on newly replicated chromatin following nuclear depletion of both Sth1 and Ino80. In this case, simultaneous depletion of both factors led to the shrinkage of most of the promoter NDRs on mature chromatin as already observed for Sth1 depletion alone, indicating that Ino80 depletion is epistatic to Sth1 depletion (fig. S4A) [23]. Thus, at steady-state, RSC can be assigned as working upstream of INO80. Read-out of the +1 nucleosome positioning on newly-replicated chromatin in the absence of these two CRs unravels a left shift indicating a dominant effect of the RSC nuclear depletion just following the replication process (Fig. 4A and B, and fig. S4B and C). Consequently, it implies that after replication, RSC is acting first, pushing the −1/+1 nucleosomes, before that INO80 restricts the expansion of NDRs.

**Fig. 4:**
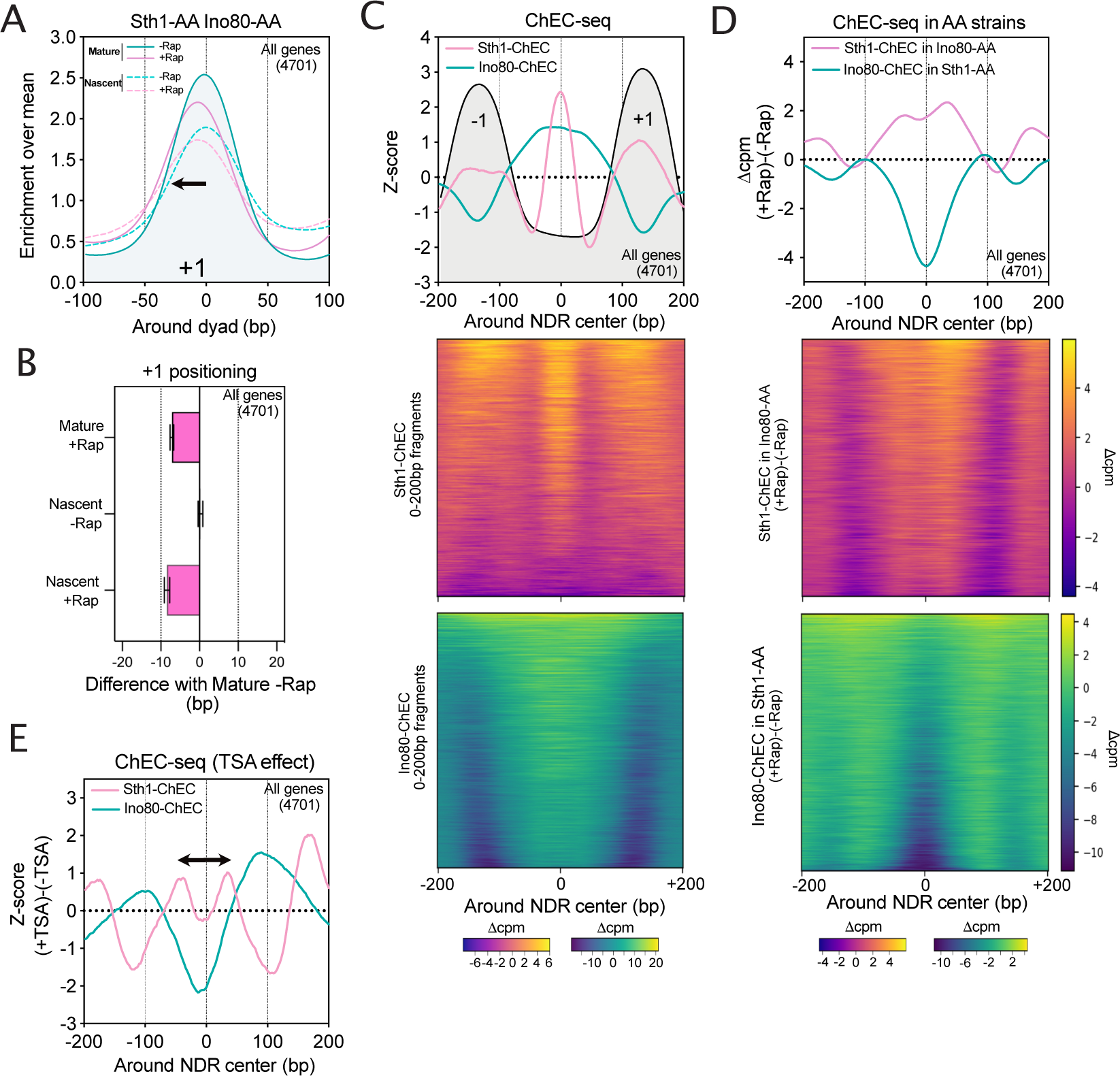
RSC acts upstream of INO80. (**A**) Metagene plot showing the mean positioning of +1 nucleosome on all genes for both ‘Mature’ and ‘Nascent’ chromatin in the presence (-Rap) or absence (+Rap) of both Sth1 and Ino80. (**B**) Bar plot showing the distribution of the difference of +1 peaks in a given condition related to the Mature -Rap position for the double Sth1-AA Ino80-AA strain. (**C**) Top panel: Average metagene plots of Sth1-ChEC and Ino80-ChEC around NDR centers. - 1/+1 positioning profiles as obtained by MNase-seq are represented in light grey. ChEC-seq data are normalized by subtracting the signal from a strain in which MNase is not fused to a given protein. Profiles are represented as a Z-score for the 0-200 bp fragments. Bottom panel: Heatmaps of the difference between Sth1-ChEC or Ino80-ChEC signals and the free MNase control. (**D**) Top panel: Average metagene plot showing the difference of Sth1-ChEC or Ino80-ChEC signals between +Rap and -Rap conditions in Ino80-AA and Sth1-AA genetic backgrounds, respectively. Bottom panel: Heatmap of the difference between +Rap and -Rap conditions for Sth1-ChEC and Ino80-ChEC in Ino80-AA and Sth1-AA genetic backgrounds, respectively. (**E**) Average metagene profile representing the difference in Sth1-ChEC and Ino80-ChEC profiles when cells are treated or not with TSA. Data are represented as Z-scores.

To confirm this result, we addressed the interplay between RSC and INO80 recruitment by ChEC-seq [50, 51]. We performed this experiment by assessing the binding of Sth1-ChEC in a Ino80-AA strain and *vice versa*. First, we considered the recruitment of Sth1 and Ino80 at steady state. As already described, Sth1-ChEC reveals a bipartite recruitment being directly in contact with the DNA at the level of the NDR center but also in contact with the neighboring - 1/+1 nucleosomes (Fig. 4C). This is in good agreement with the structural data proposing a similar conformational arrangement [48]. The profile of Ino80-ChEC shows a slightly different recruitment pattern being enriched on a broader region of the NDR and surrounding the sharp RSC/DNA contact peak (Fig. 4C). We then tackled the interdependency of RSC and INO80 (Fig. 4D). The ChEC-seq data revealed that Ino80 binding decreases for the whole set of promoter NDRs in the absence of Sth1 while the absence of Ino80 leads to a spreading of Sth1 binding around its initial peak, consistent with the view that RSC is acting upstream of INO80. Moreover, it also adds that INO80 restricts RSC positioning over NDRs. Thus, despite the need of RSC for initial recruitment of INO80, the two CRs are then entering into a “tug-of-war” as already proposed [52].

RSC is known to interact with acetylated nucleosomes *in vitro* through its multiple bromodomains and its *in vivo* recruitment at promoters is influenced by the levels of histone acetylation occurring at the surrounding −1/+1 nucleosomes [43, 58, 59]. Therefore, we tested the effect of TSA treatment on RSC and INO80 binding at promoter NDRs at steady state. Interestingly, maintenance of acetylation leads to the spreading of RSC binding being now enriched around the NDR center. This is accompanied by a shift of INO80, keeping its intercalated position between RSC contact on the DNA and the −1/+1 nucleosomes (Fig. 4E). It acknowledges a scenario in which maintenance of histone acetylation *via* TSA treatment increases the interaction with the RSC complex, hence stimulating its displacement on NDRs and setting INO80 signal accordingly.

### Steady state maturation of chromatin recapitulates the interplay between CRs and histone H3 acetylation

Our data support a scenario in which H3 acetylated nucleosomes tend to recruit RSC leading to the expansion of the NDR. Conversely, nucleosomes that are deacetylated might be better substrates for INO80.

To further test this hypothesis, we isolated extreme cases of differences in positioning between +1 peaks in nascent condition as compared to the mature one. We clustered +1 nucleosomes in two groups: the highly shifted cluster, corresponding to a right-to-left shift during maturation (>30bp), and the non-shifted cluster (Fig. 5A). When treating the cells with TSA, the highly shifted +1 nucleosome is maintained at a position similar to the nascent chromatin condition (Fig. 5A). This strongly suggests that histone acetylation reduces the pulling activity of INO80. Thus, at steady state, the +1 nucleosome of the highly shifted cluster might interact more with INO80 and become relatively deacetylated while the non-shifted cluster might be a substrate of RSC, enriched in acetylated histone H3 nucleosomes (Fig. 5B). We tested this hypothesis using the published dataset from the Friedman lab [44]. Indeed, the highly shifted cluster shows significantly lower levels of H3K9ac, H3K14ac and H3K18ac at the +1 nucleosome but not for either H3K23ac or H3K27ac (Fig. 5C). Interestingly, the most significant lower levels are observed for the H3K14ac which has been shown to be the main target of Sth1 bromodomain [53]. Moreover, the shifted cluster is significantly enriched in INO80 binding (Fig. 5D and E). Altogether, these observations are consistent with our model.

Interestingly, the highly shifted cluster is enriched in genes coding for ribosomal proteins (Fig. 5F, 4.7 fold over-enrichment, hypergeometric p-value = 1.9*10^-13^) which are already known to be INO80-dependent in terms of expression [54].

**Fig. 5:**
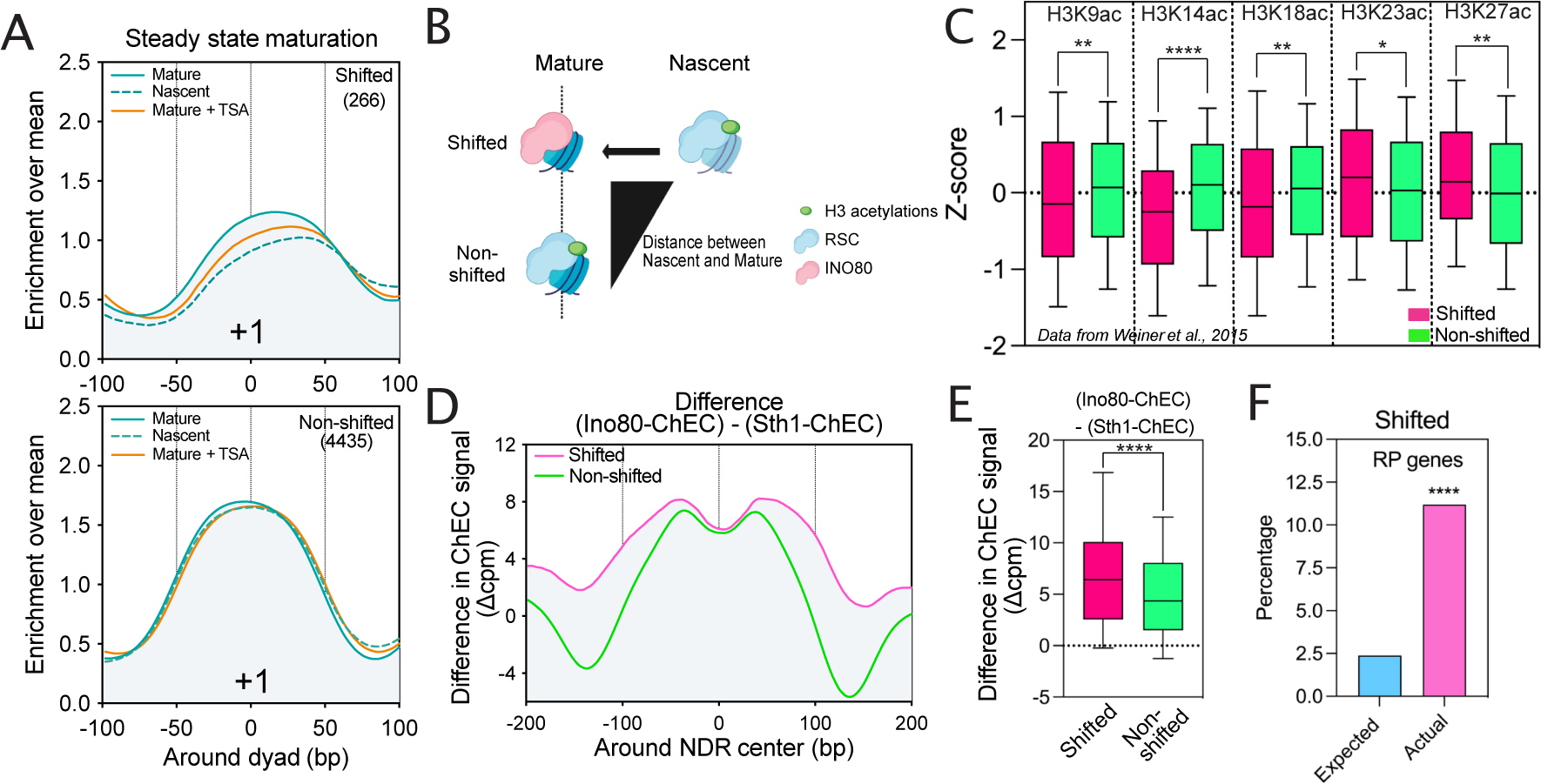
Interplay between CRs and acetylation during NDR maturation. (**A**) Average metagene plot of the +1 positioning for nascent and mature chromatin. Genes were split into two categories as ‘shifted’ and ‘non-shifted’, which were defined with the help of the DANPOS algorithm (see Materials and Methods). The metagene profile of the mature +1 positioning in cells treated with TSA is also represented. (**B**) Our model proposes that +1 nucleosomes that are shifted on nascent chromatin might be H3 acetylated, interacting with RSC and that the maturation process might decrease their H3 acetylation leading to an increased interaction with the INO80 puller. By contrast, the non-shifted ones might not decrease in H3 acetylation during the maturation process, mainly interacting with the RSC pusher complex. Thus, on mature chromatin, shifted +1 nucleosomes might be less H3 acetylated than non-shifted ones, and enriched in INO80. (**C**) Boxplot of differences in H3 acetylations on mature chromatin between shifted and non-shifted classes. Histone H3 acetylation data from Weiner et al., 2015 were normalized with their respective input by subtraction in an area of 20 bp centered on the +1 dyad and normalized as a Z-score. P-values were calculated through a Mann-Whitney test on values not normalized as Z-scores. (**D**) Average metagene profile of the difference between Ino80-ChEC and Sth1-ChEC signals for the two groups of genes aligned to the NDR centers. (**E**) Boxplot of the difference in Ino80-ChEC and Sth1-ChEC signal between Shifted and non-shifted genes. Statistics were calculated over an area of 200 bp area focused on the NDR center. P-values were calculated through a Mann-Whitney test. **(F)** Enrichment of ribosomal protein genes for the shifted cluster. The P-value is extracted from a hypergeometric distribution.

### RNAPII is not involved in promoter NDR resetting after replication

Recent results in mESCs revealed a contribution of RNAPII in resetting the accessibility of promoter NDRs after replication [36, 55]. In *S. cerevisiae*, a thermosensitive mutant of Rpb1, the catalytic subunit of RNAPII, affects the −1/+1 positioning while Rpb1 anchor-away does not reveal any striking phenotype [24, 56].

To address these discrepancies, we performed the nascent nucleosome positioning assay in the presence or absence of Rpb1. We made a distinction between the 1^st^ and 5^th^ quantiles of genes using their expression levels as a proxy to unveil their intrinsic dependence on RNAPII (Fig. 6A and B). Neither for highly nor lowly expressed genes, RNAPII appears to be important for −1/+1 positioning after replication (Fig. 6B). However, we could notice interesting differences between highly and lowly expressed genes when considering levels of +1 nucleosomes. The nascent +1 nucleosome levels of highly expressed genes are higher than those of mature chromatin in the absence of Rap. Thus, transcription restart might induce a partial depletion of +1 nucleosomes for highly expressed genes. If it is the case, +1 levels between nascent and mature chromatin in the absence of RNAPII for highly expressed genes should be equivalent, which appears to be the case. Such an effect is not observed for lowly expressed genes, pointing out the importance of transcription in +1 nucleosome depletion. Altogether, our data strongly suggest that the +1 nucleosome is first positioned after replication before being partially depleted by the transcription process implying that the transcription did not resume during the 2 min of EdU pulse. Therefore, while RNAPII may influence the mature levels of the +1 nucleosomes, the transcription process does not appear as being essential for NDR resetting post-replication in *S. cerevisiae*.

**Fig. 6:**
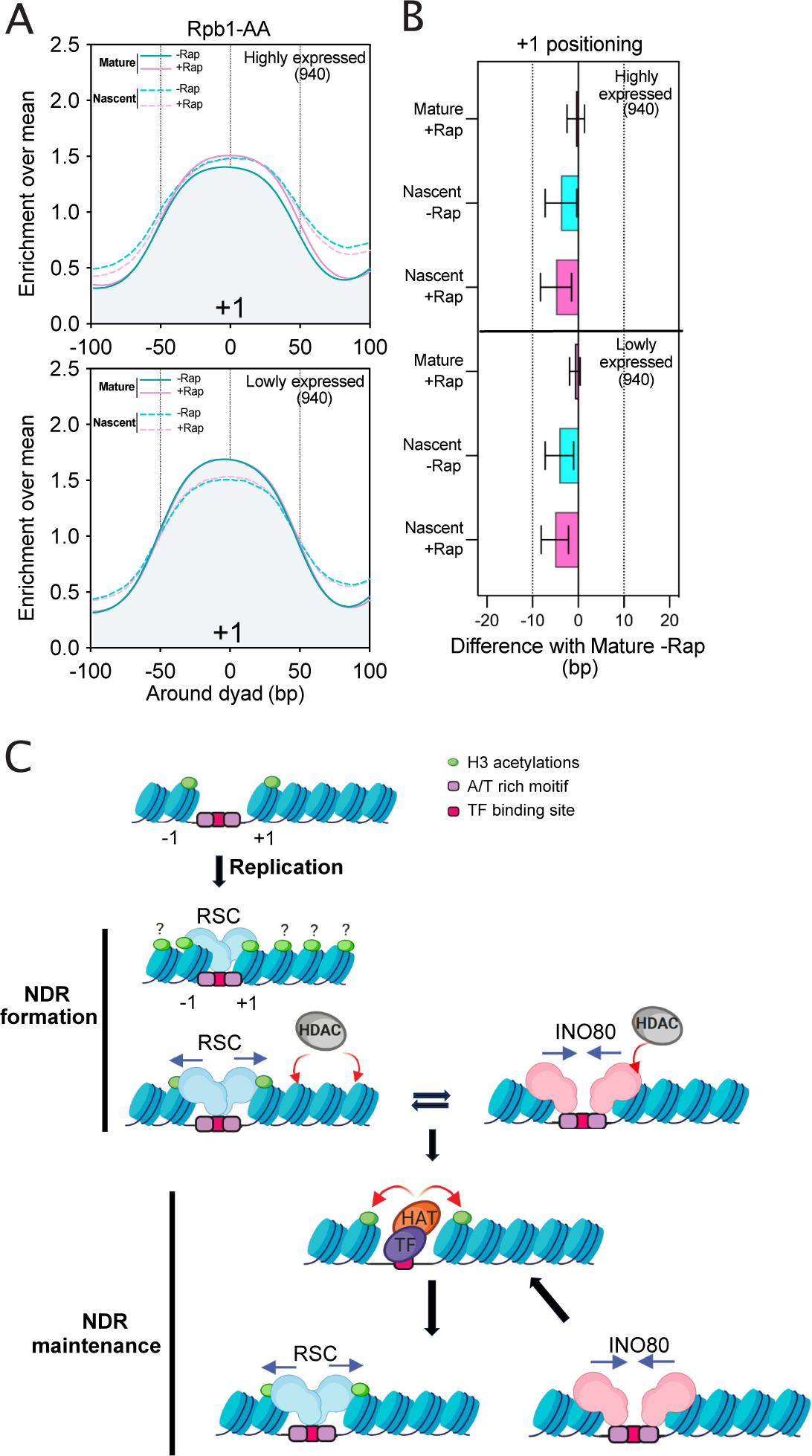
NDR reestablishment following replication is transcription-independent. (**A**) Average metagene plot of the +1 positioning for nascent and mature chromatin in the presence or absence of Rpb1. Genes were split into two categories, the top 20% and bottom 20% in terms of gene expression (as extracted from previous RNA-seq experiments, see Materials and Methods). (**B**) Bar plot showing the distribution of the difference of +1 peaks in a given condition relative to the Mature -Rap position for the Rpb1-AA strain. (**C**) Model: our data favor a two-step mechanism of promoter NDR maturation following DNA synthesis. After replication, RSC is rapidly targeted to disorganized promoters, probably *via* histone acetylation binding and/or AT-rich sequences. RSC enlarges promoter NDRs giving a window for INO80 to bind and pull the deacetylated forms of −1/+1 nucleosomes. This cycle is then maintained by TF-mediated acetylation of −1/+1 nucleosomes, favoring RSC pushing activity and so on. For simplicity, only one of the two double-stranded DNA molecules produced by replication was represented.

## DISCUSSION

Focusing on the −1/+1 positioning following replication, our work proposes a model for the sequence of molecular events leading to the resetting of promoter NDRs. First, we have shown that Reb1 and Abf1 TFs/NDFs are involved in the maintenance of NDRs but not in their reestablishment following replication (Fig. 1). We have also established that histone H3 acetylation correlates with a shift of −1/+1 nucleosomes thereby expanding promoter NDRs (Fig. 2). Interestingly, maintenance of histone acetylation rescues a proper positioning of −1/+1 in the absence of Reb1 (Fig. 2). We observed that RSC and INO80 CRs are necessary for NDR resetting post-replication with RSC acting upstream of INO80 (Fig. 3 and 4). Moreover, our data strongly suggest that RSC mainly pushes acetylated −1/+1 nucleosomes while INO80 restricts RSC activity through the pulling of deacetylated −1/+1 nucleosomes (Fig. 4 and 5). Finally, we have shown that RNAPII is not required for −1/+1 position resetting after replication (Fig. 6). Altogether, we propose a two-step model for NDR resetting and maintenance (Figure 6C).

### RSC ATPase activity as the initiator of promoter NDR resetting post-replication

How is RSC specifically targeted and/or acting at promoters? It is proposed that the sub-stochiometric subunits Rsc3 and Rsc30 constitute the DNA binding module recognizing a CGCG motif widely contained in yeast promoters [48, 49, 57]. However, our results show that Rsc3 depletion, contrary to Sth1 depletion, does not impact −1/+1 positioning immediately after replication and only affects a subset of gene promoters on mature chromatin (Fig. 3). Regarding its behavior in our assays, Rsc3 depletion phenotypes are more related to depletion of Reb1 and Abf1; Rsc3 could therefore be considered as a TF as already proposed [25, 49].

In another scenario, RSC might be recruited by acetylated histones. Indeed, RSC interacts with acetylated histones *in vitro* through its multiple bromodomains [58, 59]. Our results show that TSA treatment favors RSC pushing activity at −1/+1 nucleosomes, leading to its symmetric progression over the promoter from its nucleation site (Fig. 4). Thus, it could be appealing to propose that specific histone H3 acetylation at −1/+1 nucleosomes might trap the RSC complex post-replication. However, results obtained in yeast cells indicate that freshly deposited nucleosomes tend to be acetylated and they are not promoter specific [60]. Thus, the acetylation status might not be enough to explain this specific action at promoters. One specific feature of promoter NDRs are the poly-dA/dT stretches that induce an intrinsic depletion of nucleosomes and that are known to stimulate and orient RSC activity [13, 14, 19, 21]. Thus, the combination of AT-rich-dependent depletion of nucleosomes together with histone H3 acetylation might preferentially activate RSC at promoters and not in genes bodies.

Finally, a recent report revealed a physical interaction between RSC and the Mediator complex, working as a co-activator at promoters [61]. Whether Mediator is involved in NDR resetting after replication remains to be investigated.

### The pulling activity of INO80 follows RSC and restrains its impact

Our results clearly show that INO80 activity and recruitment are subsequent to RSC binding and pusher action (Fig. 4). Contrary to RSC, INO80 activity on chromatin is largely non-specific and is linked with longer NDRs [26]. Thus, one tempting scenario is that RSC would first push the nucleosome leaving enough linker DNA for INO80 to bind and pull the nucleosome. In agreement with such a proposal, INO80 binding is different from the RSC profile, surrounding the nucleation site of RSC. Moreover, when treating with TSA, INO80 binding is displaced in accordance with RSC binding (Fig. 4).

Another question is whether RSC and INO80 bind at the same time? This seems quite unlikely. Indeed, our results indicate that RSC activity depends on acetylated −1/+1 nucleosomes while INO80 functions more with deacetylated nucleosomes. This goes well with the “tug-of-war” model in which CRs are competing constantly to maintain NDRs [52].

Recent studies report that hexasomes are the preferred substrate of INO80 *in vitro* [62, 63]. Further studies will be needed to integrate these observations in our model.

In this study, we did not cover other CRs such as SWI/SNF or ISW2 described as acting at promoters since, unlike RSC and INO80, they only target a subpopulation of genes [23, 26].

### Reb1 and Abf1 are not acting as PFs after replication

TFs/NDFs are usually considered as equivalent to PFs in higher eukaryotes [25, 26, 64]. Indeed, Reb1 and Abf1 are involved in the maintenance of promoter NDRs *in vivo* and Reb1 interacts *in vitro* with its specific binding motif located within a nucleosome [14, 28, 29, 32]. The PF model fitted to re-organization of the chromatin after replication would place Reb1 and Abf1 as the first players in NDR resetting, which is not supported by our data thereby questioning the *in vivo* concept of PFs. Mature NDRs represent the average state of a balance between opening and closing [65, 66]. Thus, from a steady-state observation, it is impossible to extract the first occurring step. Our assay allows to follow the sequence of events and to clearly state that Reb1 and Abf1 are not acting as PFs for NDR resetting after replication.

Interestingly, TSA treatment rescues a proper positioning of −1/+1 nucleosomes in the absence of Reb1 implying that it is not the TF *per se* that maintains NDRs open, but rather their −1/+1 acetylation states (Fig. 2). Therefore, in this case, promoter NDR architecture appears to be independent of the presence of TFs. It might explain the apparent high percentage of genes in which TFs are not stably detected at yeast promoters [67]. However, our experiments did not address the transcription status in this context. Although numerous TFs show direct interactions with the transcription machinery [68], it is tempting to propose that a subset of TFs might set promoter architecture by promoting histone acetylation, which represents the keystone.

Based on this model, why would co-depletion of TFs and CRs be additive [14]? We hypothesize that the absence of TF tethering to promoters might leave enough linker DNA for INO80 to pull −1/+1 nucleosomes. Consistently, expression of the DNA-binding domain of Rap1 TF partially rescues the −1/+1 positioning [40]. Therefore, we propose that TFs have a dual function acting both as barriers limiting INO80 action and as activators of RSC pushing function by promoting histone acetylation of the flanking nucleosomes.

### Histone H3 acetylation helps the opening of promoter NDRs *via* −1/+1 nucleosomes shifts

Histones are heavily post-translationally modified leading to the hypothesis of the “histone code” [3]. While some modifications are associated with open or closed chromatin, their impact on nucleosome positioning has been poorly assessed. By reanalyzing high-resolution maps of histone modifications [44], we clearly establish that H3 acetylated nucleosomes correspond to a subpopulation of −1/+1 nucleosomes that is shifted relative to the average and associated with an expansion of the promoter NDRs (Fig. 2). In agreement with this concept, maintenance or decrease of histone acetylation shift the whole population of −1/+1 nucleosomes (Fig. 2).

Activation of genes during the metabolic cycle is associated with an expansion of promoter NDRs [69]. Strikingly, maximal expression correlates with a peak of H3K9ac and H3K18ac on +1 nucleosomes [70]. Moreover, exit from quiescence is associated with an opening of the promoter NDR as transcription resumes [71]. Altogether, it supports well our data proposing a link between H3 acetylation, chromatin opening and gene expression.

Surprisingly, acetylation of different lysine residues on histone H4 does not induce a similar shift in −1/+1 positioning despite their positive correlation with gene expression (fig. S2C) [44]. Thus, histone H4 acetylations stimulate gene expression independent of NDR expansion, probably by weakening the interaction between DNA and histones as already proposed [72].

This study, as well as our previous work, establish that H3K36me3, H3K79me3 and H4K16ac correlate with a decreased NDR width [43] (fig. S2B and C). Moreover, induction of an increase of H3K36me3 at −1/+1 nucleosomes reduces NDR length [43]. Altogether, these observations stress the importance of histone modifications in setting proper nucleosome positioning at NDRs required for optimal gene expression.

### Extension of the model to higher eukaryotes

Within the SWI/SNF family, yeast RSC is related to the BAF complexes in mammals with PBAF being the closest one [73]. All of the BAF complexes are assembled around an ATPase, being BRG1 or BRM. Very recently, a study in mESCs has shown that nearly all of the promoter NDRs, as well as enhancers, rapidly lose their accessibility when treated with an inhibitor of BRG1/BRM [74]. Importantly, this accessibility loss can be compensated with the time by the EP400/TIP60 HAT, pointing out, as in our yeast model, a probable central role of histone acetylation in promoter opening. On the other hand, in mESCs, the INO80 complex colocalizes with the TATA-binding protein (TBP) contained in the transcription pre-initiation complex marking most of the active promoters [75]. Thus, the crucial CRs of our model also vastly participate in promoter NDRs regulation in higher eukaryotes. However, their interplay with TFs and especially with PFs at promoters and enhancers remains obscure due to a lack in temporal resolution. Indeed, the two-step mechanism we propose, NDR formation and maintenance, which is involving the same players, cannot be inferred from steady state analyses. Short pulses of chromatin labeling coupled with acute depletion of TFs/CRs might reveal if this model can be extended to mammalian cells.

## MATERIALS AND METHODS

### Strains

All experiments presented in this study were performed using the budding yeast *S. cerevisiae* derived from the Anchor-Away genetic backgrounds [76]. A complete list of strains used in this study is provided in Table S1. Strains were generated by genomic integration of tagging cassettes as described [77]. Yeast cells were grown in YEP medium (1% yeast extract, 2% peptone) supplemented with 2% glucose as carbon source at 30°C. Experiments were performed with exponential phase cells harvested between an OD_600_ of 0.4 - 0.6, unless otherwise indicated. Anchor-away of targeted proteins was induced by the addition of 1 μg/mL of rapamycin to the medium for 30 min.

### Labelling of nascent DNA and crosslinking

Yeast cells from a fresh plate were grown overnight in YEPD medium at 30°C and diluted to OD_600_ = 0.3 in YEPD medium next morning. 50 mL of yeast cultures were used for each condition. After one cell cycle, rapamycin was added to corresponding samples at OD600 = 0.6 and incubated for 30 min at 30°C. For some experiments, TSA (Sigma-Aldrich, T8552) was also added for 30min at a final concentration of 1mM. Nascent DNA was labelled by addition of 5-Ethynyl-20-deoxyUridine (EdU; Roth, 7845) at a final concentration of 10 mM for 2 min. After labelling, cells were immediately fixed with formaldehyde at a final concentration of 1% for 15 minutes followed by glycine addition at a final concentration of 125 mM for 5 minutes at room temperature. Cells were washed twice with HBS (50 mM HEPES-KOH pH = 7.5, 140 mM NaCl), frozen in liquid nitrogen and kept at −80°C until further steps.

For verification of EdU pulse time, cells were arrested in G1-phase by addition of 200 ng/mL α-factor (PRIMM, 201307-00007) and incubated at 30°C for 2.5 hr. EdU was added in the medium at the end of G1 arrest for 2 min and 50 mL of G1-arrested, EdU labeled cells were crosslinked with formaldehyde. The rest of the culture was washed with YEP (no sugar) medium for three times and cells were resuspended in prewarmed YEPD medium and incubated for 30 min at 30°C to release the cells into S-phase. At the end of S-phase release, EdU was added in the medium and cells were incubated at 30°C for an additional 2min. S-phase released EdU labeled cells were then crosslinked with formaldehyde. Crosslinked cells were washed twice with HBS, frozen in liquid nitrogen and kept at −80°C until further steps.

### MNase-seq

The MNase experiments were performed mainly as described in [43] with the following modifications. For chromatin extraction, frozen cell pellets were resuspended in 600 μL cell lysis buffer (50 mM HEPES-KOH, pH = 7.5, 140 mM NaCl, 1 mM EDTA, 1% Triton X-100, 0.1% Sodium Deoxycholate, supplemented with 1 mM PMSF and EDTA-free protease inhibitors) and 600 μL of glass beads (BioSpecProducts, 11079105) was added to the cells and mechanically lysed by bead beating with a Mini-Bead beater for 1 min 30 seconds. The cell extract was then centrifuged at 13,000 rpm for 30 min at 4°C to collect chromatin. The chromatin was resuspended in 600 μL of NP buffer (0.5 mM Spermidine, 50 mM NaCl, 1mM β-mercaptoethanol, 0.075% NP-40, 10 mM Tris-HCl pH 7.4, 5 mM MgCl_2_, 1 mM CaCl_2_) per reaction. The DNA concentration was measured using Qubit dsDNA BR assay kit (Invitrogen, Q32850). Chromatin was diluted to β20-30 mg/mL using NP buffer to adjust chromatin density in each reaction. MNase treatment was performed using 0.5 μL of MNase (ThermoScientific, 88216, 100 units/mL), incubated at 37°C and 100 μL of digestion product was aliquoted for different time points (3, 6, 9 min). The MNase digestion was stopped by adding a stop buffer (20 mM EDTA, 0.5 % SDS, 0.5 mg/mL proteinase K) in the reaction and incubated first at 42°C for 1 h and then at 65°C overnight. After reversal of crosslinking, DNA was extracted with phenol-chloroform-isoamylalcohol [PCI; (25:24:1 v/v), Roth, A156], treated with RNase A (Invitrogen, 12091021) and visualized on a 2.5% agarose gel. The nucleosomal DNA sample with the best MNase digestion time point (corresponding roughly to a ratio of 2 for mononucleosomes/dinucleosomes) was purified again with PCI extraction before proceeding further steps.

### Click reaction and streptavidin affinity capture

MNase digested and purified DNA fragments were subjected to a click reaction that adds biotin to the EdU-labeled newly synthesized DNA. Click reaction was performed in the following order to the indicated final concentrations and protected from light: 25 ng of DNA was mixed with 1 mM Biotin-PEG-Azide (Sigma-Aldrich, QBD10825). In a separate tube CuSO_4_ (Sigma-Aldrich, C1297) and THPTA (Sigma-Aldrich, 762342) were added at 2 mM/4 mM ratio, respectively and mixed by vortexing (CuSO_4_/THPTA mix, 1/2 ratio). Then DNA/Biotin PEG Azide mix was combined with CuSO4/THPTA and 10 mM ascorbic acid (Roth, 3525) was added to the mix. The reaction final volume was adjusted to 50 μL by using 10 mM Tris-HCl (pH = 7.5), mixed by vortexing and incubated at 37°C for β 2 hours with shaking at 800 rpm.

The click reaction product was purified by ethanol precipitation. Biotin conjugated EdU-labeled DNA was then pulled down using 5 μL of Dynabeads M-280 streptavidin (Invitrogen, 11205D). Beads were pre-washed three times with 1x Tween buffer (5 mM Tris HCl pH = 8, 0.5 mM EDTA, 1M NaCl, 0.05% tween) before being resuspended in 2x B&W Buffer (10 mM Tris HCl pH = 8, 1 mM EDTA, 2M NaCl) and combined with an equal volume of biotinylated DNA. The reaction was incubated at room temperature for 30 min with gentle shaking. The beads were washed twice with 1x Tween buffer and once with 1x B&W Buffer (5 mM Tris HCl pH = 8, 0.5 mM EDTA, 1M NaCl). Following washing steps, bead-bound DNA was eluted by adding sterile water and heating at 70°C for 5 min.

### ChEC-seq

ChEC-seq experiments were performed mainly as described in [43] with the following modifications. The yeast strains were cultured the day before in YEPD medium and next morning they were diluted to OD_600_ = 0.2 in YEPD. The 30 min rapamycin treatment (1 μg/ mL, final) was performed for anchoring away of targeted proteins and/or 30 min TSA treatment (1 mM final) at OD_600_ = 0.5. For each condition, 50 mL cell cultures were harvested at room temperature. The cells were washed once in 1 mL Buffer A (15 mM Tris-HCl pH 7.5, 80 mM KCl, 0.1 mM EGTA, 0.2 mM spermine, 0.5 mM spermidine and EDTA-free protease inhibitor tablet (Roche, 04693159001). Cells were resuspended in 693 μL of buffer A and 7 μL of 10% digitonin (0.1% final concentration; Sigma-Aldrich, D141) were added to facilitate permeabilization of the cells for 5 min at 30 °C and mixed with the pipette several times during the incubation. CaCl_2_ was added (final concentration 5 mM) to activate the MNase cleavage. For each sample 100 μL of cells were collected at 30 s after the induction of MNase digestion and were immediately mixed with 200 mL of 2X Stop solution (400 mM NaCl, 20 mM EDTA, 4 mM EGTA, 1% SDS). The cells were treated with Proteinase K (0.4 mg/mL final concentration) and incubated at 55 °C for 30 min. The DNA was extracted using PCI extraction and precipitated with EtOH by adding 30 μg glycogen. Following RNase treatment, size selection of DNA fragments was performed as follows; 100 μL of AmpureXP beads (Beckman Coulter, A63881) were added in 40 μL of RNase-treated DNA and placed on a magnetic rack. The supernatant was then transferred to the new tube containing 10 mM Tris and 0.2 M NaCl for each reaction. The DNA was extracted with PCI and ethanol precipitated. The purified DNA samples were then used for library preparation using NEBNext UltraII DNA Library prep kit (NEB, E7370L) and were sequenced at the iGE3 platform of the University of Geneva.

### Data analyses

For MNase-seq experiments, fastq files were first trimmed from their adapters using Trim Galore! (https://github.com/FelixKrueger/TrimGalore). Then, trimmed files were aligned using Bowtie2 on the sacCer3 genome in a paired-end manner with the following options: ‘-k 20– end-to-end–sensitive −X 800’ [78]. BigWig files were generated using bamCoverage from deepTools2 [79] (with the following options: normalization in counts per million (cpm), bin size of 1bp, exclusion of the chrM for normalization, regions centred with respect to the fragment length, a minimum mapping quality of 20, duplicates removal, size selection of 120 to 200 bp. For ChEC-experiments, the same processing was followed with the exception of a size selection from 0 to 200 bp. Means of experiments were obtained with the bigwigCompare command from deepTools2 with a bin size of 1bp. Differences in signal between conditions were also calculated using bigwigCompare. Metagene plots and heatmaps were obtained using the plotProfile and plotHeatmap commands from deepTools2, respectively. Enrichment over mean plots were made as followed: average plots were calculated for the whole set of −1/+1 nucleosomes in the −100 to +100 bp window around the dyad, then every position relative to the feature was divided by the average number of reads over this −100/+100 bp window. It allows to normalize the profiles of nascent chromatin showing a reduced read depth due to purification. For most of the experiments, the means of two replicates are presented in main figures while separate replicates are depicted in supplementary figures. For Fig. 1, 2, 3, 4 and 6, the difference in +1 peak position per gene was calculated using a custom Python script deposited on Zenodo. For Fig. 5, the shifted and non-shifted populations were extracted using DANPOS3 (https://github.com/sklasfeld/DANPOS3) with the replicate 2 of the Reb1-AA strain in the absence of Rapamycin.

### -1/+1 annotations and gene/replication origins lists

All the coordinates and gene/replication origins names are listed in Table S2. −1/+1 coordinates and NDR centers were obtained from [43] and [80], respectively. Replication origins coordinates were retrieved from [81]. Reb1 targets were extracted from the RNA-seq analysis in a Reb1-AA strain from [40]. Abf1 targets were retrieved from the Rpb1 ChIP-seq in an Abf1-AA strain from [42]. Rsc3 targets were defined from a k-means clustering of the difference in MNase-seq profile around +1 nucleosome in the Rsc3-AA (Fig. 3). Highly transcribed and lowly transcribed genes were retrieved from the RNA-seq in an Nrd1-AA strain in the absence of Rap [41].

### Reanalysis of existing data

Data from [45] for MNase-seq in absence or presence of TSA or with the *epl1* mutant were analyzed following the same procedure as the data analyses section. For MNase-(ChIP)-seq data from [44], single end reads were virtually extended to 150 bp and the centers of these fragments were extracted using deepTools2. For each of the histone modification presented, the corresponding total MNase-seq signal (Input) was subtracted.

### Statistical analyses and model design

Plots and statistical analyses of this work were performed using Prism 10.0 (Graphpad). All tests are nonpaired tests. Mann–Whitney *U* tests were used to extract a *P*-value. ns if *P*-value > 0.05, ^∗^ < 0.05, ^∗∗^ < 0.01, ^∗∗∗^ < 0.001, ^∗∗∗∗^ < 0.0001. Models were designed using BioRender.

## Acknowledgements

We thank Audrey Noireterre and Guillaume Canal for critical reading of the manuscript, comments and discussions. We are grateful to Geraldine Silvano for technical assistance and for the good mood of the lab. We thank José Manuel Nunes for many discussions about nucleosome analyses. We also thank the iGE3 genomics platform for sequencing our samples in a timely manner.

## Author contributions

Conceptualization: SZ and JS. Methodology: SZ, JKG and JS. Investigation: SZ, JKG and JS. Data analysis: JS. Writing-original draft: JS. Writing-review and editing: SZ, FS and JS. Funding acquisition: FS.

## Data and materials availability

The accession number for the data reported in this study is GEO: GSE262171. The custom python script used to find +1 peaks was deposited on Zenodo *via* https://zenodo.org/records/10866695.

---

**Figure S1 – related to Fig. 1:**
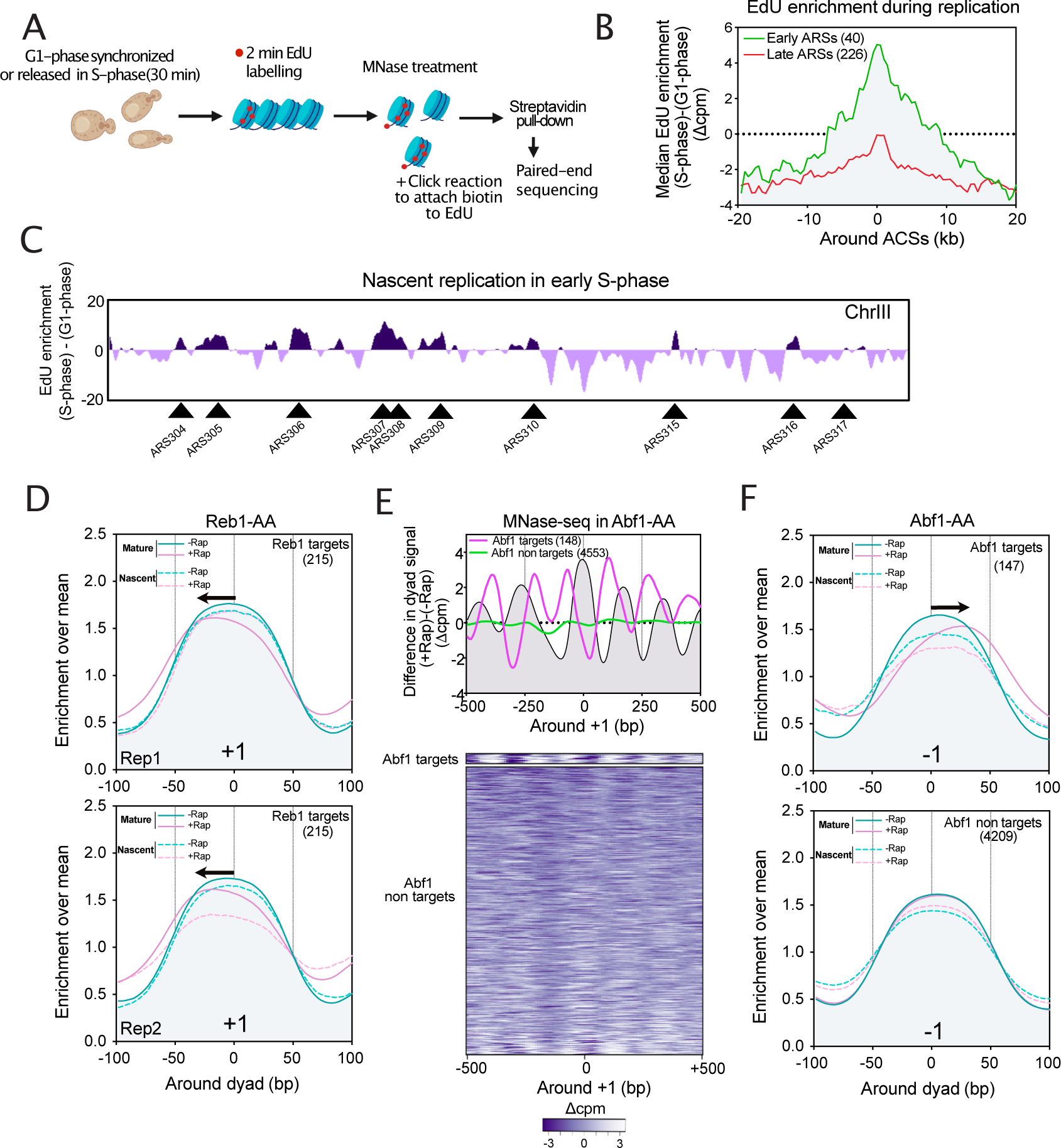
(**A**) Schematic representation of the experimental design to define EdU labeling efficiency for purification of nascent chromatin: Cells were synchronized in G1-phase for 2.5 hr before being released in S-phase for 30min at 30°C. EdU was added in the medium for 2min during G1 arrest and at 30min after S-phase release. Cells were then crosslinked, the chromatin was extracted and treated with MNase. After a click reaction to attach biotin, newly synthesized DNA was purified with streptavidin beads. The nascent DNA was finally paired-end sequenced. (**B**) Metagene plot representing the median of the difference between S- and G1-phase cells. Replication origins were sorted as two classes: Early ARSs and Late ARSs, based on [81]. (**C**) Difference between S- and G1-phase for the whole chromosome III. Early ARSs are indicated by arrows. (**D**) Same as Fig. 1C for Reb1 targets at +1 for each of the two replicates. (**E**) Same as Fig. 1B for Abf1-AA. (**F**) Same as Fig. 1E at −1 nucleosome.

**Figure S2 – related to Fig. 2:**
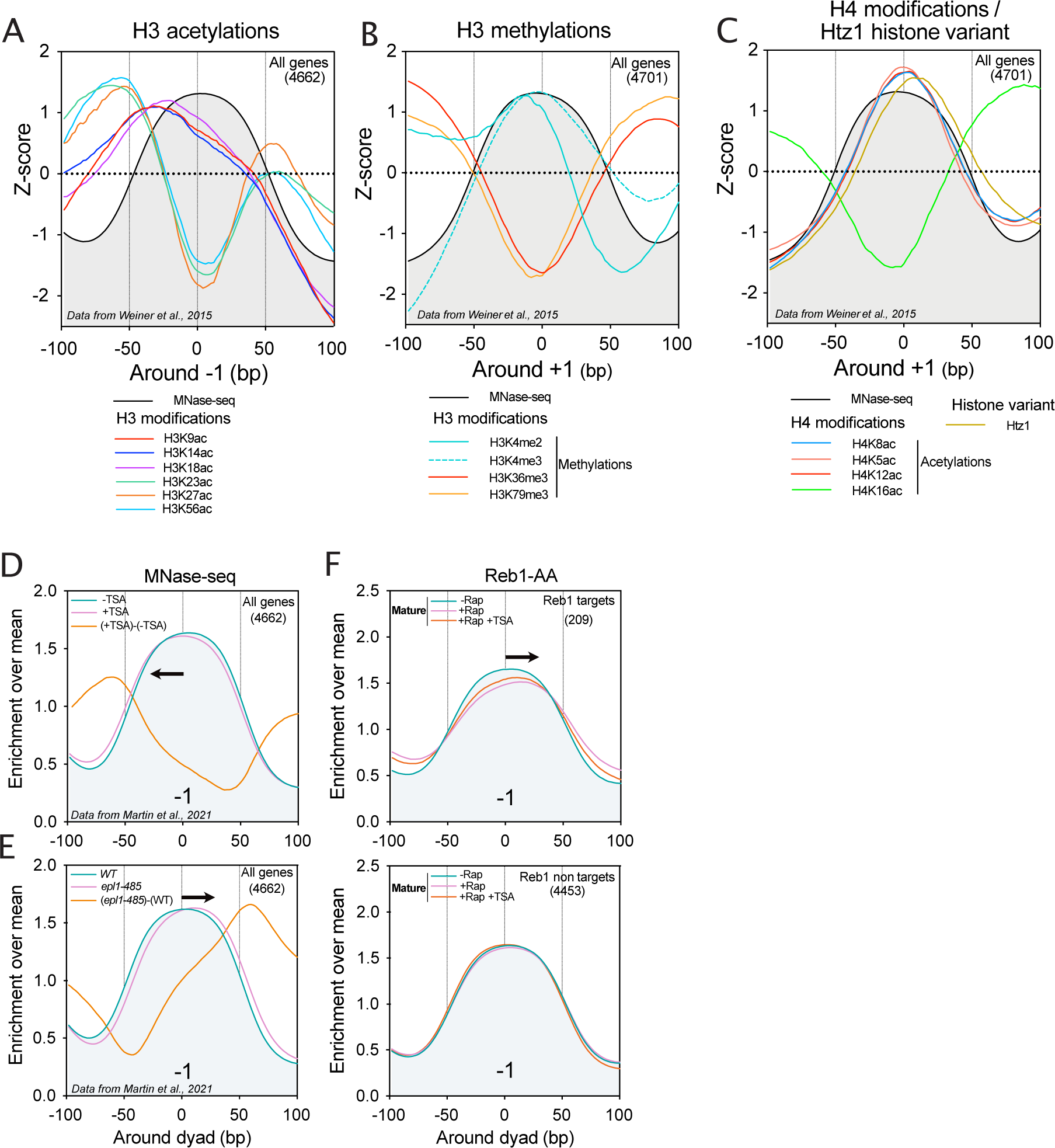
(**A**)(**B**)(**C**) Same as Fig. 2B, respectively, for acetylation of different lysine residues on histone H3 around the −1 dyad, for methylation of different lysine residues on histone H3 around the +1 dyad and for acetylation of different lysine residues on histone H4 around the +1 dyad using data from Weiner et al., 2015. (**D**)(**E**)(**F**) Same as Fig. 2C, E and F, respectively, for the −1 nucleosome.

**Figure S3-related to Fig. 3:**
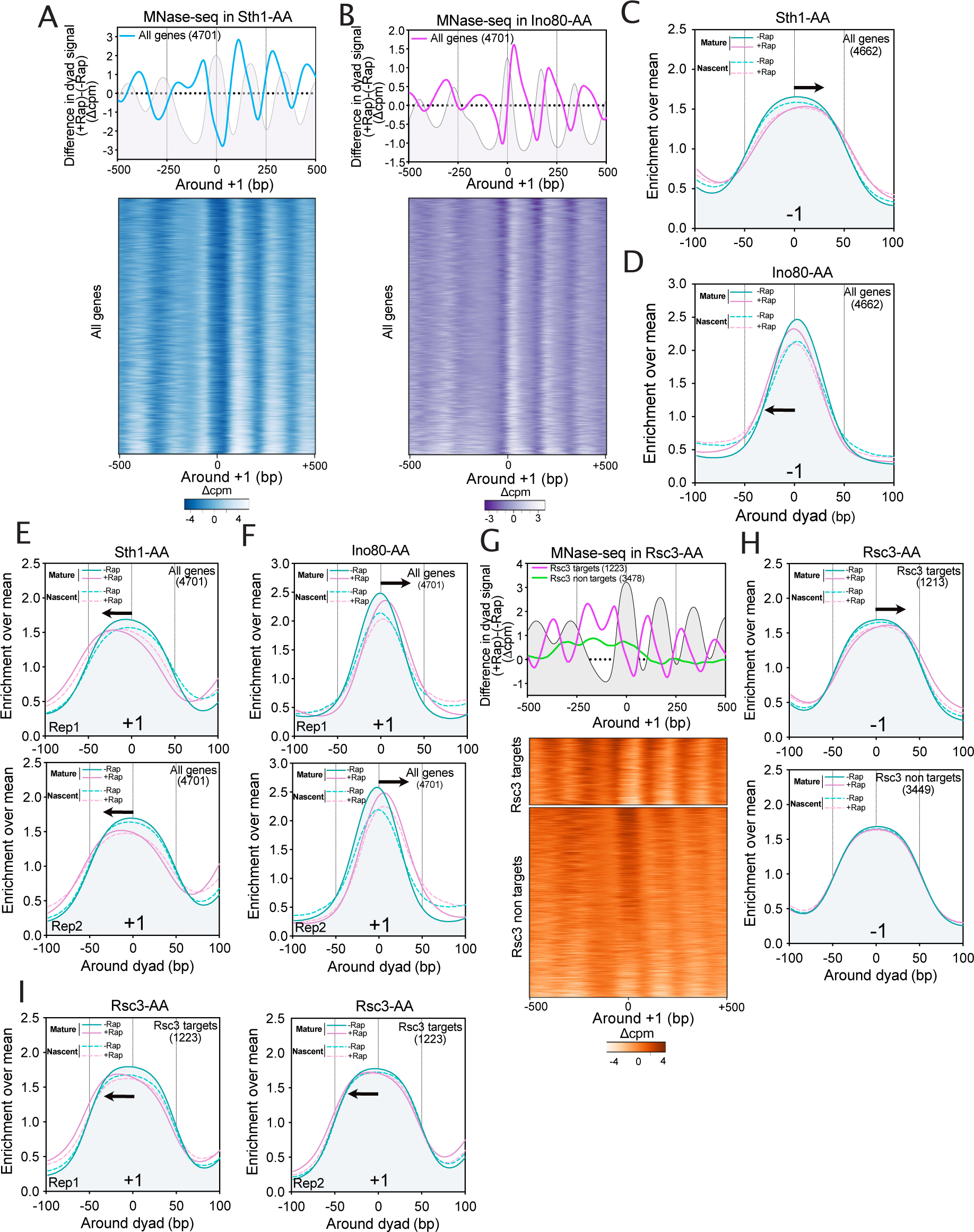
(**A**) (**B**) Average metagene plot and heatmaps of the difference in MNase-seq profiles for Sth1-AA (A) and Ino80-AA (B) strains. The grey profile represents the MNase-seq signal in the absence of Rap, in order to visualize the position of −1/+1 nucleosomes. (**C**) (**D**) Same as Fig. 3, A and C for the −1 nucleosome. (**E**) (**F**) Same as Fig. 3, A and C for each replicate. (**G**) Average metagene plot and heatmaps of the difference in MNase-seq profiles for the Rsc3-AA strain. (**H**) Same as Fig. 3E for the −1 nucleosome at Rsc3 targets. (**I**) Same as Fig. 3E for replicates at Rsc3 targets.

**Figure S4-related to Fig. 4:**
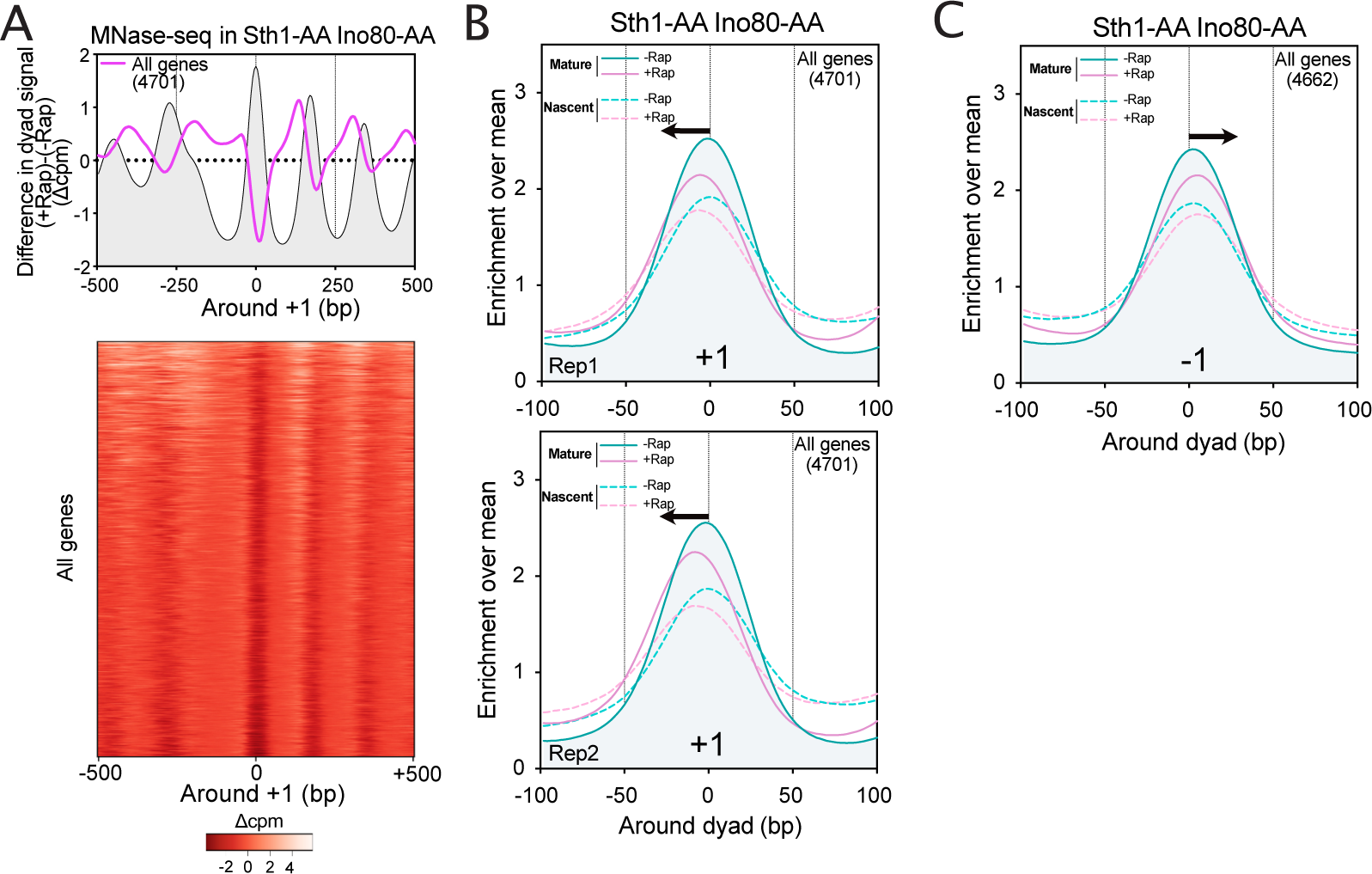
(**A**) Average metagene plot and heatmaps of the difference in MNase-seq profiles for the double Sth1-AA Ino80-AA strain. (**B**) Same as Fig. 4A for each of the two replicates at +1 nucleosome. (**C**) Same as Fig. 4A for the −1 nucleosome.

**Table S1:** Yeast strains used in the study.

| name | genotype | Source | Figure |
| --- | --- | --- | --- |
| FSY8613 | HHY221 (MATa tor1-1 fpr1::loxP-LEU2-loxP RPL13A-2×FKBP12::loxP ade 2-1 trp1-1 can1-100) dBAR1::URA3 BrdU::HIS3(HIS+) | Haruki et al., 2008 | Fig. S1A-C |
| FSY8614 | HHY221 (MATa tor1-1 fpr1::loxP-LEU2-loxP RPL13A-2×FKBP12::loxP ade 2-1 trp1-1 can1-100) dBAR1::URA3 BrdU::HIS3(HIS+) Reb1-FRB::KAN | This study | Fig. 1B-D; Fig. 2F-G; Fig. S1D; Fig. S2F |
| FSY8615 | HHY221 (MATa tor1-1 fpr1::loxP-LEU2-loxP RPL13A-2×FKBP12::loxP ade 2-1 trp1-1 can1-100) dBAR1::URA3 BrdU::HIS3(HIS+) Rsc3-FRB::KAN | This study | Fig. 3E-F, Fig. S3G-I |
| FSY8618 | HHY221 (MATa tor1-1 fpr1::loxP-LEU2-loxP RPL13A-2×FKBP12::loxP ade 2-1 trp1-1 can1-100) dBAR1::URA3 BrdU::HIS3(HIS+) Abf1-FRB::KAN | This study | Fig. 1E-F; Fig. S1E-F |
| FSY8789 | HHY221 (MATa tor1-1 fpr1::loxP-LEU2-loxP RPL13A-2×FKBP12::loxP ade 2-1 trp1-1 can1-100) dBAR1::URA3 BrdU::HIS3(HIS+) Rpo21-FRB::KAN | This study | Fig. 6A-B |
| FSY9654 | HHY221 (MATa tor1-1 fpr1::loxP-LEU2-loxP RPL13A-2×FKBP12::loxP ade 2-1 trp1-1 can1-100) dBAR1::URA3 BrdU::HIS3(HIS+) Sth1-FRB::KAN | This study | Fig. 3A, B, G, H; Fig. S3A, C, E |
| FSY 9587 | HHY221 (MATa tor1-1 fpr1::loxP-LEU2-loxP RPL13A-2×FKBP12::loxP ade 2-1 trp1-1 can1-100) dBAR1::URA3 BrdU::HIS3(HIS+) Sth1-FRB::KAN INO80-FRB::NAT | This study | Fig. 4A-B, Fig. S4 |
| FSY 9377 | HHY221 (MATa tor1-1 fpr1::loxP-LEU2-loxP RPL13A-2×FKBP12::loxP ade 2-1 trp1-1 can1-100) dBAR1::URA3 BrdU::HIS3(HIS+) Sth1-Flag-MNase::HYG | This study | Fig. 4C, E; Fig. 5D-E |
| FSY 9384 | HHY221 (MATa tor1-1 fpr1::loxP-LEU2-loxP RPL13A-2×FKBP12::loxP ade 2-1 trp1-1 can1-100) dBAR1::URA3 BrdU::HIS3(HIS+) Sth1-FRB::KAN INO80-MNase::HYG | This study | Fig. 4C-E; Fig. 5D-E |
| FSY9452 | HHY221 (MATa tor1-1 fpr1::loxP-LEU2-loxP RPL13A-2×FKBP12::loxP ade 2-1 trp1-1 can1-100) dBAR1::URA3 BrdU::HIS3(HIS+) INO80-FRB::KAN | This study | Fig. 3C-D, G-H; Fig. S3D, F |
| FSY9456 | HHY221 (MATa tor1-1 fpr1::loxP-LEU2-loxP RPL13A-2×FKBP12::loxP ade 2-1 trp1-1 can1-100) dBAR1::URA3 BrdU::HIS3(HIS+) INO80-FRB::KAN Sth1-Flag-MNase::HYG | This study | Fig. 4D |
| FSY8295 | W303 RAD5+ MAT::loxP ADE2::loxP LEU2::pGAL-HO LYS2::loxP pREB1-MNase-3xFLAG- URA3 dBAR1::HIS3 | This study | Fig. 4C |

